# The effect of combined dexmedetomidine and isoflurane anaesthesia on visually evoked responses in rats – an electrophysiological study

**DOI:** 10.1101/2022.07.05.498786

**Authors:** Freja Gam Østergaard, Christian Stald Skoven, Bettina Laursen, Tim B. Dyrby, Alex Wade

**Affiliations:** H. Lundbeck A/S, Ottiliavej 9, 2500 Valby, Denmark; Danish Research Centre for Magnetic Resonance, Centre for Functional and Diagnostic Imaging and Research, Copenhagen University Hospital - Amager and Hvidovre, Copenhagen, Denmark; Department of Psychology, The University of York, Heslington, York, YO10 5DD, United Kingdom; Department of Applied Mathematics and Computer Science, Technical University of Denmark, Kongens Lyngby, Denmark

**Author notes:** University of Groningen, Groningen Institute for Evolutionary Life Sciences, Groningen, Netherlands (Freja Gam Østergaard). Center for Functional Integrative Neuroscience (CFIN), Aarhus University, Aarhus, Denmark. Corresponding author’s.

## Abstract

The purpose of this study was to pilot an anaesthetic regime with the potential to be used for blood oxygenation level dependent functional magnetic resonance imaging (BOLD fMRI) of the visual system. We used electrophysiology to explore the effect of an anaesthesia regime, combining dexmedetomidine and isoflurane, on the visual system of female Sprague-Dawley rats. This paradigm is hypothesised to affect neural signalling less than other paradigms, and thus may be suitable for studies using BOLD fMRI. Electrodes were implanted bilaterally in the visual cortex (VC) and superior colliculus (SC) of 16 rats. Visual evoked potentials (VEPs) and steady-state VEPs (SSVEPs) were recorded during 1 Hz and 14 Hz light stimulation, respectively. As a control experiment an exploration of the wash-out of isoflurane was performed. The combined anaesthetic regime showed, that the visually evoked responses were almost completely abolished at the initiation of anaesthesia, but gradually recovered to baseline levels over time in the SC. In contrast, the evoked response in the VC only partially recovered, staying significantly below baseline condition. BOLD fMRI results from a previously published study supports this finding.

## Introduction

Preclinical blood oxygenation level dependent (BOLD) functional magnetic resonance imaging (fMRI) is traditionally carried out in anaesthetized animals. Anaesthesia in general is known to impact both the neural signalling and the neurovascular coupling that underlies the BOLD fMRI signal (Desai et al., 2011; Gao et al., 2017). Isoflurane (iso), which is a very commonly used anaesthetic, can attenuate the BOLD signal due to vasodilation and increases in cerebral blood flow (Lee et al., 1994). However, the combination of the anaesthetic dexmedetomidine (dex), with a low dose of isoflurane has previously been suggested as suitable for BOLD fMRI (Benveniste et al., 2017; Magnuson et al., 2014; Pan et al., 2013). Dexmedetomidine is an α2-agonist with considerable vasoconstrictive effects (Weerink et al., 2017), possibly counteracting some of the effects of isoflurane.

This electrophysiological study was carried out prior to an fMRI experiment (Østergaard et al., 2021) to assess the effect of an anaesthetic protocol combining dexmedetomidine with a low dose of isoflurane (dex/iso) on neuronal activity. We used the rodent visual pathways as a model system and assessed visual evoked potentials (VEP) and steady-state VEP (SSVEP) with the goal of optimizing the anaesthesia regime to obtain reliable a BOLD fMRI signal. In principle the regime should work similarly with other sensory modalities. The combined anaesthesia was compared to a pure isoflurane condition. Furthermore, the wash-out of isoflurane was assessed to explore any lingering effects influencing neuronal activity.

This study shows that dex/iso provides a stable anaesthesia with the possibility of detecting event-related responses, making it suitable for BOLD-fMRI. Further, we have included an fMRI scan obtained along with the scans in Østergaard et al. 2021, showing that measurable BOLD responses can indeed be obtained under dex/iso anaesthesia. As the wash-out of iso shows substantial lingering effects, this could explain why the dex concentration needs to be increased an hour after reduction of isoflurane, and why it takes three hours before the evoked potential is stable.

## Methods

### Animals

This study was carried out at H. Lundbeck AS, Copenhagen (DK) in accordance with the European Communities Council Directive (86/609/EEC), and in accordance with Danish law on care of laboratory animals. The experimental protocol was approved by the Danish Animal Experiments Inspectorate (Forsøgsdyrstilsynet) prior to the initiation of the study.

16 female Sprague-Dawley rats underwent stereotactic surgery where six electrodes were implanted. The electrodes were placed bilaterally in visual cortex (AP: −6 mm, ML: ±4 mm), and superior colliculus (AP: −6 mm, ML: ±1 mm, DV: −3.5 mm). A reference electrode (AP: +8 mm, ML: −2 mm) and a ground electrode (AP: −2, ML: +4) were also implanted. Coordinates were respective to bregma (for AP), the sagittal suture (for ML) and dura (for DV) - and were guided by (Paxinos and Watson, 1998). The animals were the same as the ones used in the study (Østergaard et al., 2020). Both the surgical details and the stimulation paradigms have been published in that article. Four animals were excluded from the present study due to poor data quality.

### Anaesthesia

Anaesthesia was induced at time 0 min, using isoflurane 5%, and a tail-vein catheter was placed. As soon as the catheter was in place, dexmedetomidine (dexdormitor, Orion Pharma, DK) was infused at 0.05 mg/kg/hr (Pump 11, Pico Plus Elite, Harvard Apparatus, US), and isoflurane turned down to 0.5% (in 1 L/min of oxygen:medical air, 8:2). Isoflurane at 0.5% was continued throughout the experiment. Blood oxygen level were monitored throughout the experiment using a pulse-oximeter (Nonin BV., NL).

Published data guided the choice of dexmedetomidine dose and infusion timing (see table 1). At time 60 min, the infusion of dexmedetomidine was increased to 0.1 mg/kg/hr. This concentration is lower than suggested by the literature, as the SpO2 measurement was destabilized after an hour with the higher concentration, likely due to vasoconstrictive effects of dexmedetomidine (Weerink et al., 2017).

**Table 1.** Comparison of anaesthetic regimes used in the literature with visual stimulation. (IV) intravenous; (IM) intramuscular; (SC) subcutaneous; (SC) superior colliculus; (VC) visual cortex; (dLGN) dorsal lateral geniculate nucleus.

| Induction |  | Maintenance |  | Article | Stimuli | Anatomical region activated |
| --- | --- | --- | --- | --- | --- | --- |
| isoflurane | 4% | urethane | 1.3g/kg bolus (IP),<br>0.13g/kg supplements (IV) | (Bailey et al., 2013) | RGB LEDs,<br>optical cables | SC+VC+dLGN |
| isoflurane | 2.5% | medetomidine,<br>pancuronium<br>bromide | 0.1mg/kg/h<br>2mg/kg/h | (Pawela et al., 2008) | 5ms, 465nm<br>(blue) LED | SC+VC+dLGN |
| ketamine,<br>xylazine | 75 mg/kg<br>5 mg/kg | medetomidine | Bolus 0.08mg/kg □<br>0.15mg/kg/h (IM) | (Van Camp et al., 2006) | Strobe unit<br>(bi-, monocular) | SC+VC |
| isoflurane | 4% | isoflurane | 1% | (Lau et al., 2011b, 2011a) | Green LED 10Hz, Fibre optics | SC+dLGN+(VC) |
| isoflurane | 2% | medetomidine | Bolus 0.04mg/kg<br>0.08mg/kg (SC) | (Niranjan et al., 2016) | cold white LED | SC+VC+dLGN |
| isoflurane | 5%<br>1.5-2% | medetomidine | Bolus 0.1mg/kg<br>0.2mg/kg /h (SC) | (Pradier et al., 2021) | Unilateral blue laser (488 nm) | SC+VC+dLGN |
| isoflurane | 5%<br>2.5% | isoflurane<br>dexmedetomidine | 0.5%<br>0.05-0.1 mg/kg/h (IV) | This study and<br>(Østergaard et al., 2021) | White LED, fibre optics | SC+(VC+dLGN) |

### Experimental design

The stimulation paradigm consisted of a 1 Hz on/off continuous flickering of 20 lux warm white LED for 400 s for 1 Hz VEPs and followed by a faster steady-state paradigm (SSVEP) at 14 Hz for 100 s and was the same as used in Østergaard et at 2020. The light was delivered from LEDs 40 cm above the animal, lighting up the entire cage. Lux measurements were made at the bottom of the cage. Electrophysiological recordings of local field potentials (LFPs) were performed in freely moving animals.

#### Isoflurane condition

recordings were made before induction of anaesthesia (time: −10 min), t: +30 min after induction and every ten minutes until t: +120 min after induction. Isoflurane anaesthesia was turned off after the recording at t: +30 min, as the VEP varies with the concentration of isoflurane used (data not shown)(Magnuson et al., 2014) and the recordings were discontinued at t: +120 min.

#### Dex/iso condition

isoflurane was reduced to 0.5 % at t: 0 min, and the infusion of dexmedetomidine was initiated. Recordings were made at time −10 min, t: +30 min and every 10 min until t: +180 min.

All animals underwent both anaesthetic paradigms with at least two weeks between recording sessions. The stimulus paradigm and recording were controlled using Spike2 v7.20 (Cambridge Electronic Design Ltd, UK). The recorded signals were sampled at 1 kHz, amplified and filtered (low-pass filter: 200 Hz; high-pass filter: 1 Hz) online using a Brownlee amplifier model 410 (Brownlee Precision, CA, US).

One BOLD-fMRI data set was obtained from one animal, that was a part of the study (Østergaard et al., 2021), where the fMRI method and data processing is described. Briefly, the animal was anaesthetised similarly to the dex/iso condition: anaesthesia was induced with iso (5% 0.5%), and dex infusion was initiated at 0.05mg/kg/h, then the animal was placed in the scanning bed of a 7T Bruker Biospec preclinical scanner, and a surface receive coil was taped to its head. Echoplanar images were acquired at t: +120 min and +180 min, with a slice thickness of 0.5 mm, TR = 1500 ms, and TE = 8.35 ms.

### Data processing and statistical analysis of VEP

The EEG signal was averaged in Spike2 yielding the VEP. Latency from stimulation to VEP peak and amplitude from baseline to VEP peak were extracted manually (peak designations are shown in Figure S1, after the reference list) and exported to R for statistical analysis. The electrophysiological signal was analysed with 48 one-way ANOVAs (7 peaks from VC + 5 peaks from SC, both amplitude and latency for pure iso condition and combined dex/iso (7+5)*2*2=48), testing for the influence of time from induction (baseline condition), on the latency and amplitude of the individual peaks. The p-values were adjusted for multiple comparisons using false discovery rate (Benjamini and Yekutieli, 2001). The p-values are shown in Table 2 for the comparisons where p<0.05.

**Table 2.** Table of p-values from the 48 one-way ANOVAs from the quantifications of VEPs. All results refer to differences relative to the baseline condition. The mean and SD of each peak can be found in table S1 (after references).

| Adjusted p-values from one-way ANOVAs |  |  | Isoflurane ☐ awake<br>n=12 |  |  | Isoflurane/dexmedetomidine<br>n=12 |  |  |
| --- | --- | --- | --- | --- | --- | --- | --- | --- |
|  |  |  | +30 min | +120 min | +180 min | +30 min | +120 min | +180 min |
| Superior colliculus | Amplitude | P1 |  |  |  |  |  |  |
|  |  | N1 | 0.02 |  |  |  |  |  |
|  |  | P2 | 0.00965 |  |  |  |  |  |
|  |  | N2 | 0.0289 |  |  |  |  | 0.00476 |
|  |  | P3 |  |  |  |  |  |  |
|  | Latency | P1 | 0.0101 |  |  | 9.31E-06 | 0.00251 | 0.00227 |
|  |  | N1 |  |  |  | 0.00361 |  | 0.0406 |
|  |  | P2 |  |  |  | 0.00776 |  |  |
|  |  | N2 |  |  |  | 0.042 |  |  |
|  |  | P3 |  |  |  | 0.00159 | 0.000775 |  |
| Visual cortex | Amplitude | P1 |  |  |  |  | 0.0152 | 0.000499 |
|  |  | N1 |  |  |  | 2.15E-05 |  |  |
|  |  | P2 |  |  |  |  | 0.00848 | 3.56E-07 |
|  |  | N2 | 0.0398 |  |  |  |  | 0.00713 |
|  |  | P3 | 0.000585 |  | 7.66E-05 |  |  |  |
|  |  | N3 | 0.000748 |  |  | 2.90E-12 | 0.00262 |  |
|  |  | P4 |  |  |  | 0.000241 |  |  |
|  | Latency | P1 |  |  |  | 1.81E-05 | 6.16E-09 | 1.03E-06 |
|  |  | N1 |  |  |  | 0.0412 | 0.00632 |  |
|  |  | P2 |  |  |  | 0.00175 | 0.00234 |  |
|  |  | N2 |  |  |  | 0.00127 | 3.15E-05 | 0.01058 |
|  |  | P3 |  |  |  | 0.0013 | 7.31E-05 |  |
|  |  | N3 | 0.00239 |  |  | 0.000193 | 0.00119 |  |
|  |  | P4 |  |  |  |  | 1.86E-05 | 8.86E-05 |

### Data processing and statistical analysis of SSVEP

The SSVEP data was separated from the VEP data and converted to mat files in Spike2. These files were imported into Matlab (Mathworks, MA, US) and time-frequency spectra were produced using part of Accusleep’s source code (Barger et al., 2019). Further analysis were carried out using the Rstudio interface for R (Rstudio, MA, US) and the R package DABEST for estimation statistics (Ho et al., 2019). Here 5000 bootstrap resamples of the data were carried out to estimate the true difference of means.

## Results & Discussion

### Visual evoked potentials (VEPs)

The purpose of this pilot was to study the effect of anaesthesia on visual responses measured as local field potentials during exposure to light flashing at a constant rate.

The results from the VEP assay of both paradigms are shown in Figure 1 for the VC, and Figure 2 for the SC. The results from one-way ANOVAs are shown in the table 2. To keep and animal anaesthetised with isoflurane the concentration needs to be adjusted regularly throughout the recording session, the concentration ranged between 1.5-2.5%. The shape of the VEP show large variation which seemed to correlate with the concentration of isoflurane. Figure 1 and 2 show that the VEP waveforms are close to being normalized at t: +120 min, i.e., approximately 90 minutes after the cessation of isoflurane. In the dex/iso condition (Figure 1, right panels), the recordings continued for three hours. The shapes of the VEPs are similar to the baseline condition at +180 min, but with minor yet significant changes in the peaks of the VEP (see table 2, means and SD as listed in table S1, after the references).

**Figure 1.**
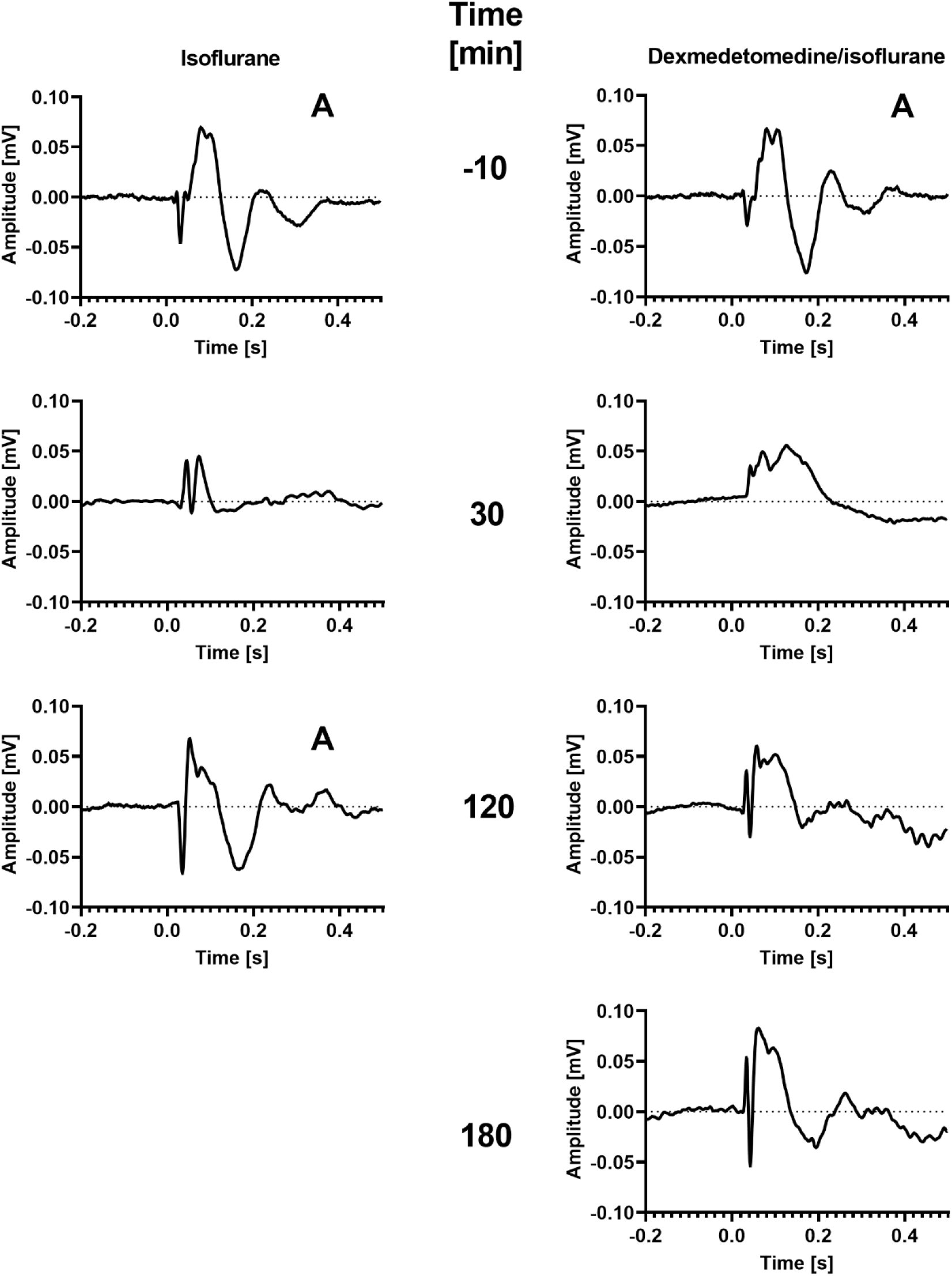
Anaesthesia affects the VEP in the visual cortex. The numbers between the columns refer to time since initiation of anaesthesia. For the isoflurane condition, the animals were only anaesthetized at time 30 min (the EEG response varied with the level of isoflurane). Right after the recording obtained at this timepoint, the anaesthesia was ceased. At time 120 min, the animals had been awake for 60-70 minutes. In the dex/iso condition, the animals were anesthetized throughout the recording session, and anaesthesia was ceased after the recording at 180 minutes. ‘A’ marks recordings obtained during the awake state. The peak designations are shown in the supplement.

**Figure 2.**
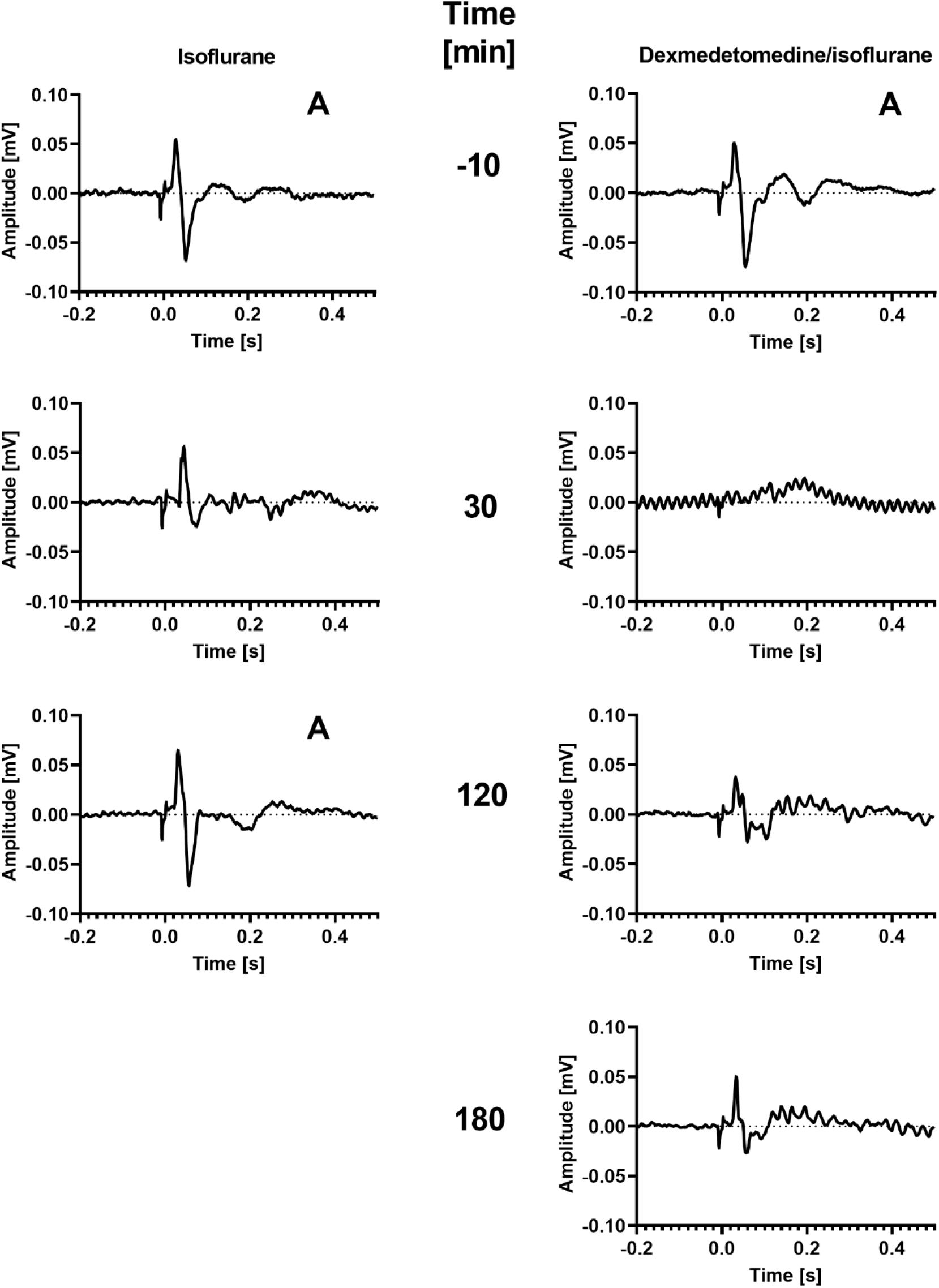
Anaesthesia affects the VEP recorded in the superior colliculus. The numbers between the columns refer to time in minutes since initiation of anaesthesia. ‘A’ marks the recordings performed when the animal was awake.

For the dex/iso paradigm, table 2 shows that the number of peaks that differ significantly from baseline, decreases over time. At t: +180 the VEP was almost normalized in both the VC and the SC. However, even after three hours there are still some changes relative to baseline.

### Steady-state visual evoked potential (SSVEP)

Figure 3 shows time-frequency plots for the concatenated SSVEP data averaged across 12 animals. The recordings for each spectrogram, corresponding to a timepoint, were 100 s long and recorded every ten minutes. The animals were only anaesthetized with isoflurane during the 30 minutes time span (form t: 0 min to t: 30 min). However, a general tendency to sleep was noticed after cessation of isoflurane. Sleep and active states were recognized from the power band. Sleep was seen most prominently in the response from the visual cortex, as the high-powered low-frequency bands at time 40 and 70-90 min. In contrast to the sleep state, the active state was characterized by a power increase in the frequency band around 8 Hz (low alfa) and one at <4 Hz (delta), as seen at times −10 and 100-120 min. The line at 14 Hz is the direct response to the flickering light stimulus, with the second harmonic of the response visible at 28 Hz. The visual cortex seemed to be affected more than the SC. Suggesting that isoflurane has a differentiated effect on cortical neurons compared to deeper neurons.

**Figure 3.**
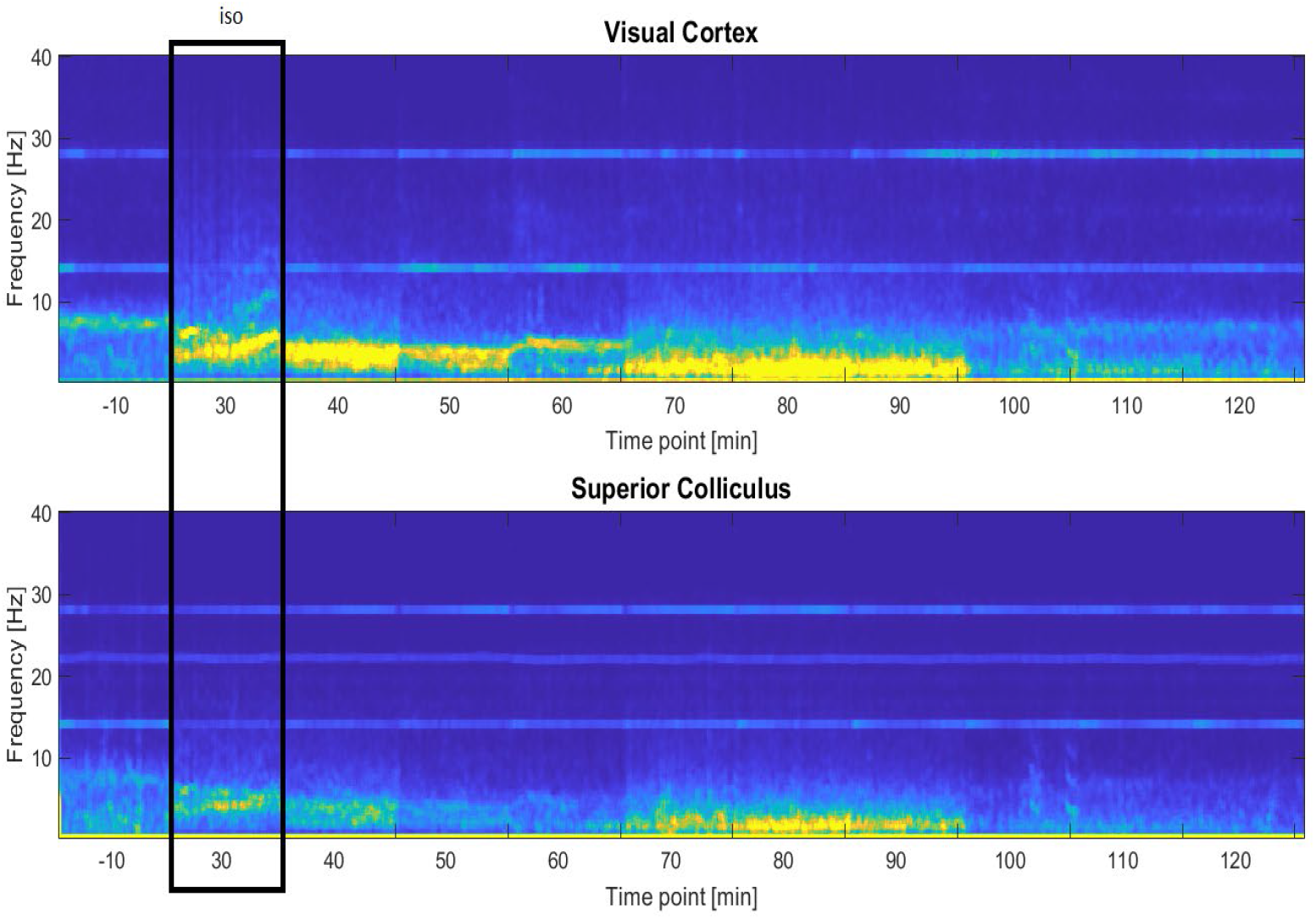
Grand average time-frequency spectra of SSVEP responses during and after isoflurane anaesthesia. The top panel shows the recordings from the visual cortex. The bottom panel shows the response from the SC. The SSVEP recording lasted for 100s with light flickering at 14 Hz and was repeated every 10 minutes. The animals were only under isoflurane anaesthesia at timepoint t: 30 min. At this point, the power of SSVEPs seemed to correlate directly with the level of isoflurane causing very large inter-individual variation. There is a clear response at 14 Hz along with the first harmonic at 28 Hz. In the SC, the band at 22 Hz was caused by a heating pad which was replaced during the course of the experiment. The superior colliculus seemed to be less affected by isoflurane than the cortex.

The main purpose of this pilot was to characterize the effects of combined anaesthesia dex/iso on neural activity. Figure 4 shows the time-frequency plot for the dex/iso condition. The animals were anaesthetized from t: 0 min and onwards. The general response demonstrated high power in the delta band (0.5-4 Hz), similar to what is seen during natural sleep (Benveniste et al., 2017). The visual response seemed to be abolished immediately after the induction of anaesthesia. The response was, however, recovered over time (Table 3). The lower panels of Figure 4 show the power in the bands at 14 Hz and 28 Hz for VC and SC normalized to baseline. The response from SC appears to be completely rescued at t: +120 min, while the response in the VC is only partially recovered. Maintaining anaesthesia using iso alone was difficult and required constant small adjustments of the iso concentration. The combined iso/dex did not require any adjustments of the concentration of iso. The concentration of dex was doubled at t: +60 min, but no further adjustments of concentration was required.

**Figure 4.**
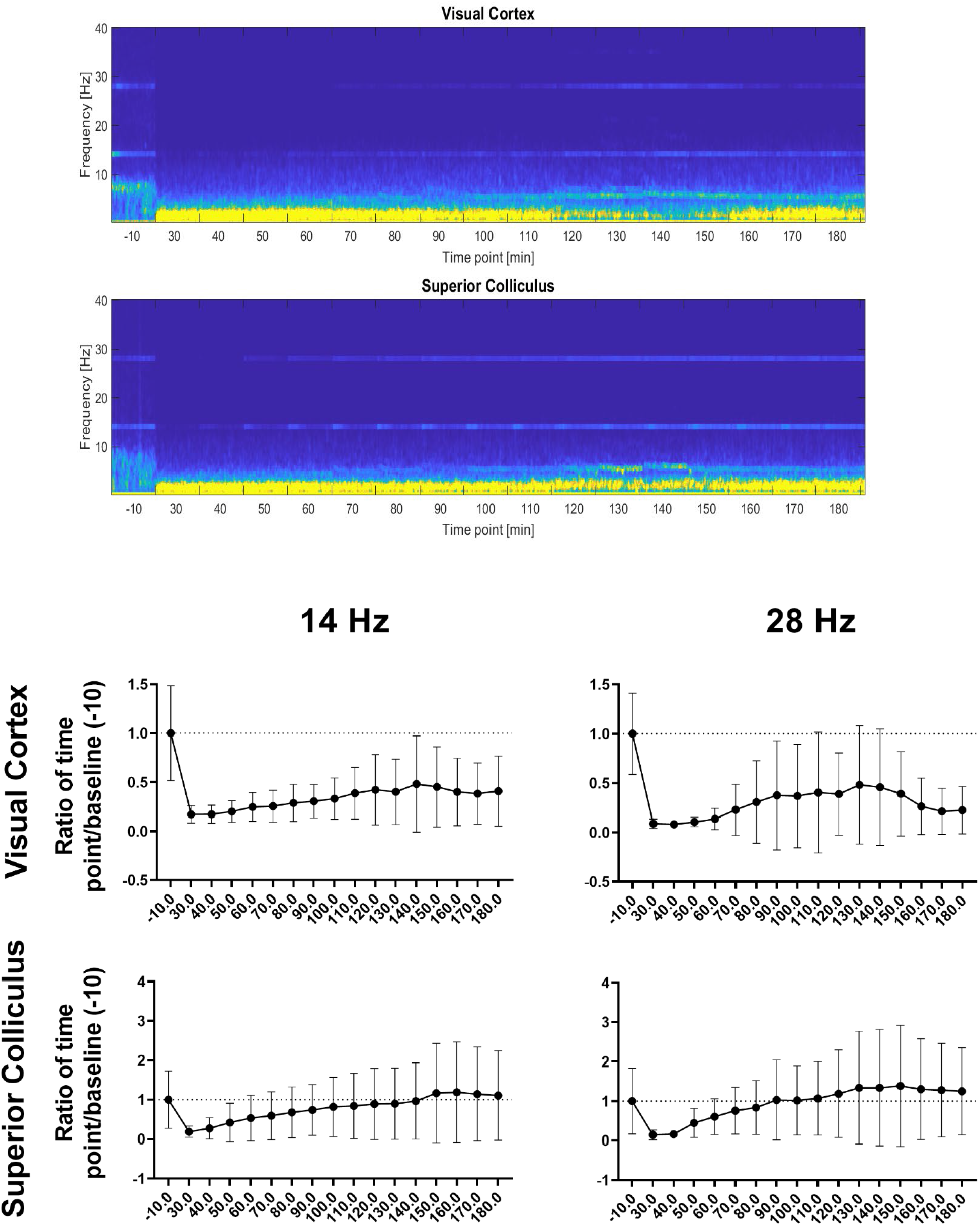
SSVEP response under anaesthesia with dexmedetomidine and low dose isoflurane. Top panels show time-frequency spectra of the SSVEP response. This response seems to be completely abolished in both the VC and SC at the induction of anaesthesia, but over the course of three hours, the responses recovered completely in the SC, and to some extend in the VC. Bottom panels: Power of SSVEP extracted from time-frequency plots (above). The power is depicted as ratio of timepoint over baseline (timepoint −10) with mean±SD over subjects. This suggests that the VC is more affected than the superior colliculus. The average of the SC seems to overshoot after 130-150 minutes, but with a large SD.

**Table 3.**
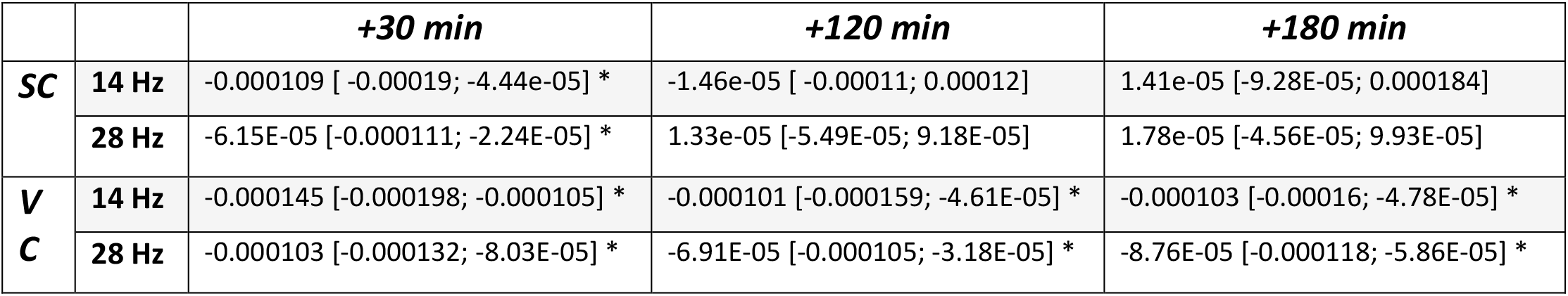
Estimation statistics of powerbands shown with difference of means compared to baseline and 95 % confidence interval of the difference of mean. Based on 5000 bootstrap resamples. Asterisks indicate that the difference of means is significantly different from zero.

|  |  | **+30 min** | **+120 min** | **+180 min** |
| --- | --- | --- | --- | --- |
| **SC** | **14 Hz** | -0.000109 [-0.00019; -4.44e-05] * | -1.46e-05 [-0.00011; 0.00012] | 1.41e-05 [-9.28E-05; 0.000184] |
|  | **28 Hz** | -6.15E-05 [-0.000111; -2.24E-05] * | 1.33e-05 [-5.49E-05; 9.18E-05] | 1.78e-05 [-4.56E-05; 9.93E-05] |
| **V** | **14 Hz** | -0.000145 [-0.000198; -0.000105] * | -0.000101 [-0.000159; -4.61E-05] * | -0.000103 [-0.00016; -4.78E-05] * |
| **C** | **28 Hz** | -0.000103 [-0.000132; -8.03E-05] * | -6.91E-05 [-0.000105; -3.18E-05] * | -8.76E-05 [-0.000118; -5.86E-05] * |

The estimation statistics for power at three timepoints (t:+30min, +120min and +120min) relative to baseline are shown in table 3. Only the power in the SC returned to baseline over the course of the experiment. The responses obtained from the VC were significantly different from baseline throughout the recording sessions. This effect may be related to effect shown in the iso condition: that the cortical neurons are affected more by anaesthesia than the subcortical.

These results show that dex/iso provides a stable anaesthesia with the possibility of detecting event-related responses. Further, we have included a fMRI scan from Østergaard et al. 2021 as an example (figure 5), showing that measurable BOLD can indeed be obtained under dex/iso anaesthesia (the specific scans have not been published before).

**Figure 5.**
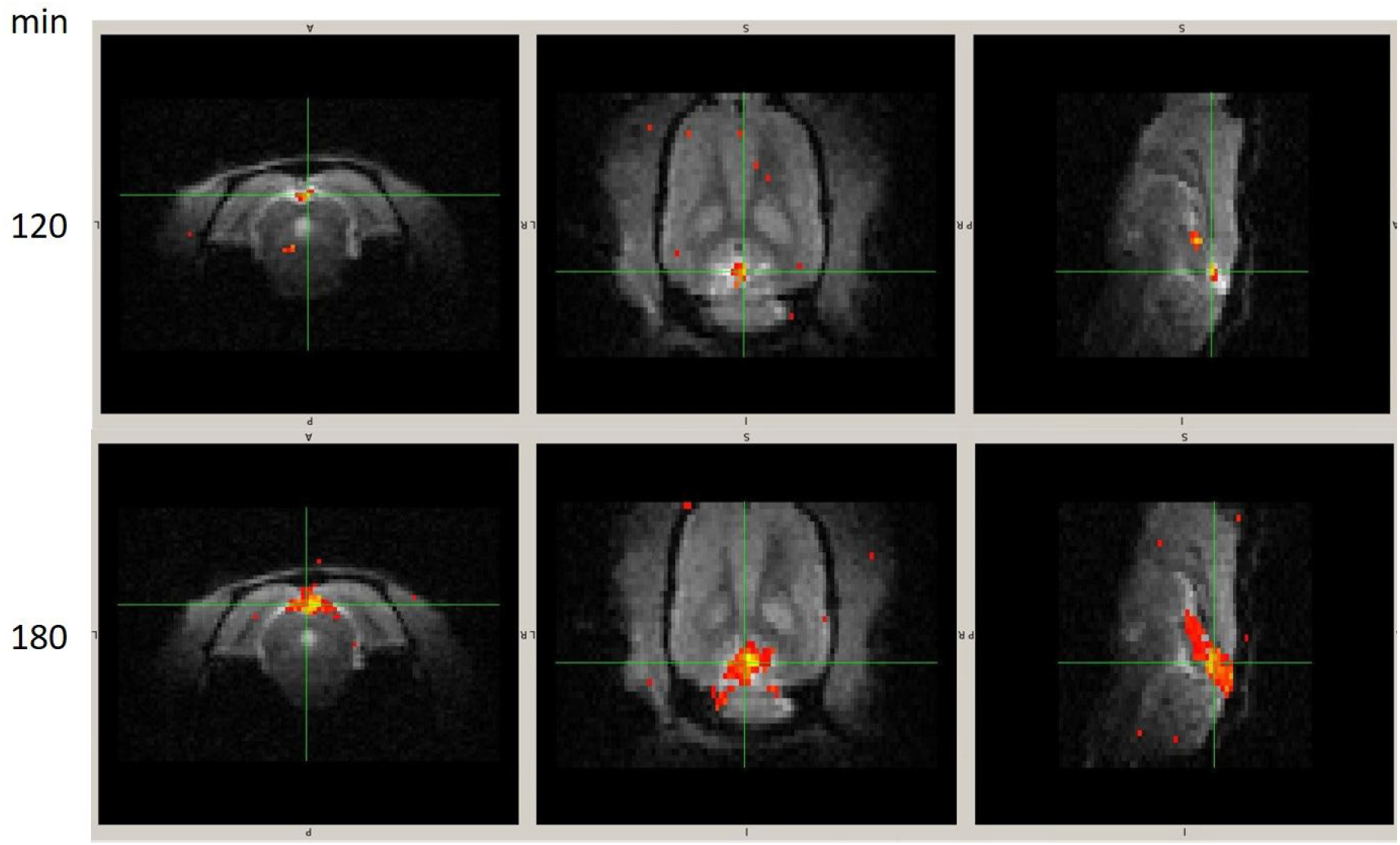
Effect of anaesthetic on BOLD (n=1). The first row shows the BOLD 120 minutes after the dexmedetomidine infusion has been started. The second row shows the same animal after 180. There is no spatial smoothing and the threshold is the same for both panels (min=3.5 SD from zero).

Figure 5 shows BOLD-fMRI from one rat under dex/iso anaesthesia at t: +120 min and t: +180 min. Here, the number of active voxels (BOLD > 3.5 SD from 0) in the SC seemed to be increased at t: +180 min, compared to t: +120 min. This is consistent with the results shown above. The BOLD response is most likely a summation of the response and all harmonics (Engell et al., 2012). As shown in the 2021 paper by Østergaard et al., activity in the VC was still detectable after 180 min. This suggests a partial correlation between power and BOLD response - the power at time 120 min and time 180 min is roughly the same, but the number of active voxels increases, indicating that the hemodynamic response is more sensitive to the lingering effect of isoflurane than the neuronal response. The hemodynamic response depends on the neurovascular coupling, which is a slower response, as it requires changes in activity in larger populations of neurons (Huneau et al., 2015) than what is required for LFPs.

Apart from testing the isoflurane and dex/iso conditions, a recording session was also carried out using hypnorm/midazolam (Figure S2) to study whether this anaesthetic would have a potential. This fentanyl-based anaesthetic largely abolished the visual response. Additionally, this anaesthetic has another practical downside, as it precipitates after approximately two hours at room temperature.

## Conclusion

In the dex/iso condition, stable VEPs and SSVEPs were obtained at t: +180 min (3 hours) after decreasing isoflurane to 0.5 %, and the anaesthetic regime described allows for use in longitudinal experimental designs.

## Summary

The purpose of the study was to study an anaesthetic regime with the potential to be used for BOLD fMRI. Isoflurane alone caused a considerable change in the VEP (Figures 1 & 2, Table 2 and Table S1) compared to the un-anaesthetized state. During the recording sessions, we found that the VEP response varied with the level of isoflurane indicating a direct dose-response relationship (Figure 3), and lingering effects on the VEP after cessation. To overcome both the reduction and instability of signal introduced by isoflurane, a combination of low-concentration isoflurane and the infusion anaesthetic dexmedetomidine (Pawela et al., 2009) was employed. This combination demonstrated minimal impact on the neural signal in agreement with other studies (Benveniste et al., 2017; Magnuson et al., 2014). One apparent downside of this regime is that it takes 3 hours (Figures 4) from the isoflurane is reduced, until a stable fMRI signal can be obtained (Figure 5). This could be a result of lingering effects of isoflurane (Magnuson et al., 2014) affecting neurovascular coupling.

## Acknowledgements

FGØ and ARW were supported by the European Union’s Horizon 2020 research and innovation programme under the Marie Skłodowska-Curie Grant Agreement No. 641805. The experimental work was carried out as part of FGØs PhD project at the Neurograd PhD school of the University of Copenhagen. Hartwig Roman Siebner acted as main faculty supervisor, participated in the conceptional planning of the project, supervision at project meetings, and have provided critical input to the interpretation of the results and was involved in drafting of the manuscript.

## Author contributions

FGØ, TBD, CSS: Conception of the project idea and project planning; FGØ, CSS: carried out the experiments, FGØ: data analysis and statistics, interpretation of results and the first manuscript draft. All authors contributed to the manuscript writing.

## Supplementary

**Figure S1.**
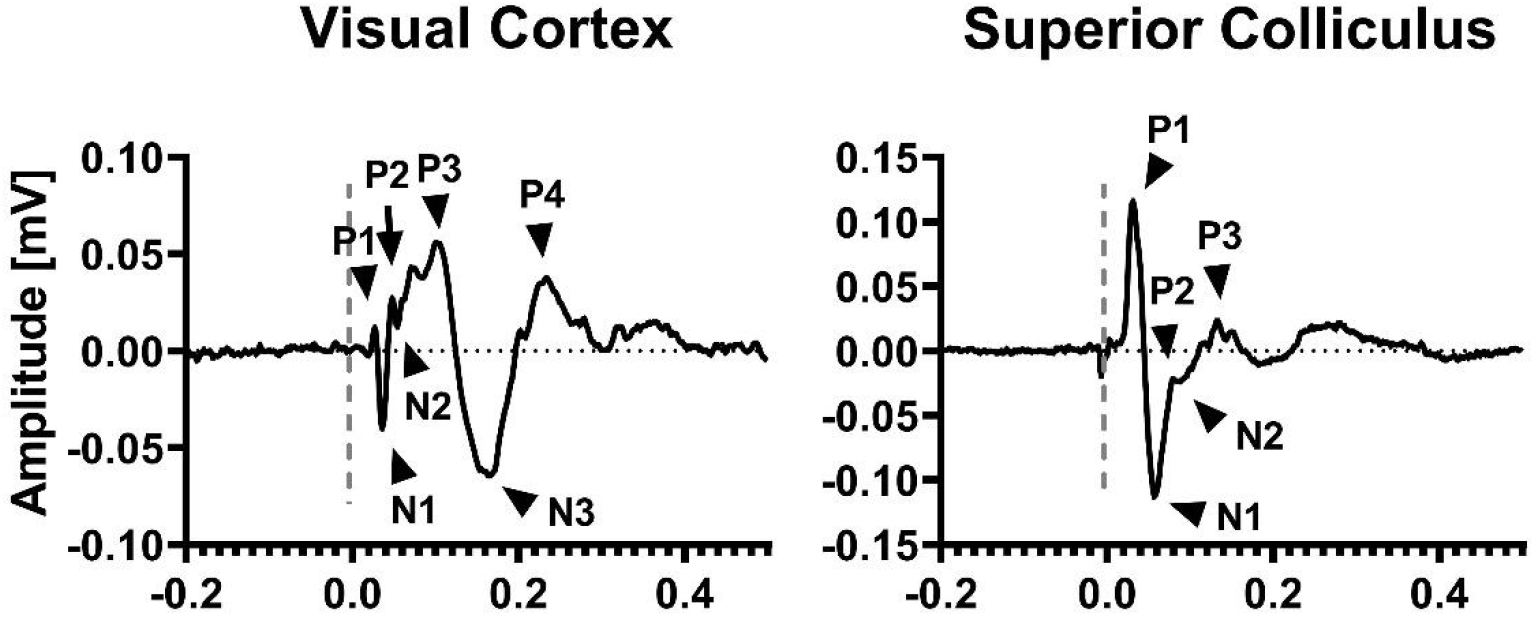
Peak definitions for VEPs recorded in the visual cortex and superior colliculus

**Table S1.**
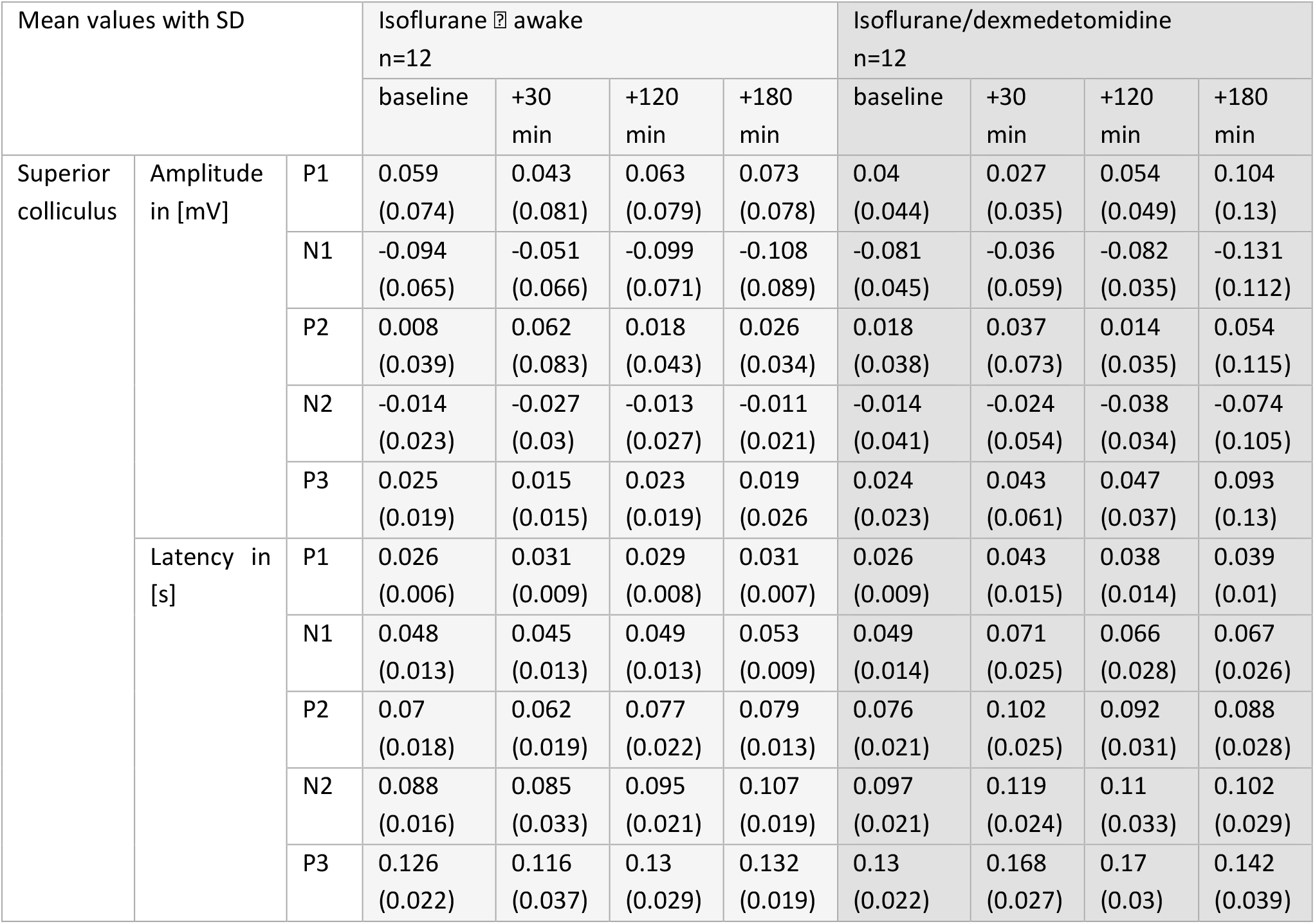
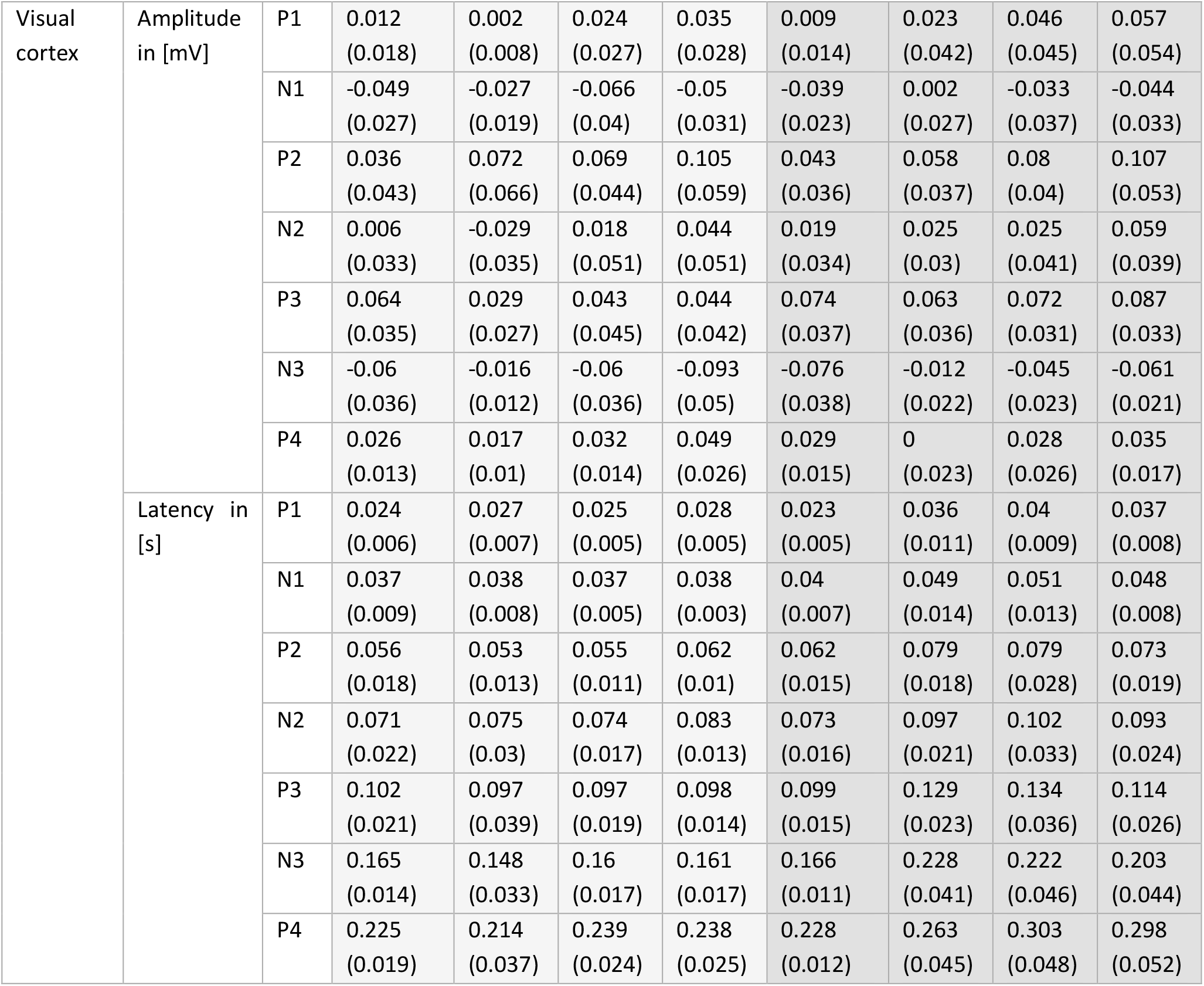
Mean and standard deviation (SD) of amplitudes and latencies from VEP waveforms

| Mean values with SD |  |  | Isoflurane ☐ awake<br>n=12 |  |  |  | Isoflurane/dexmedetomidine<br>n=12 |  |  |  |
| --- | --- | --- | --- | --- | --- | --- | --- | --- | --- | --- |
|  |  |  | baseline | +30<br>min | +120<br>min | +180<br>min | baseline | +30<br>min | +120<br>min | +180<br>min |
| Superior<br>colliculus | Amplitude<br>in [mV] | P1 | 0.059<br>(0.074) | 0.043<br>(0.081) | 0.063<br>(0.079) | 0.073<br>(0.078) | 0.04<br>(0.044) | 0.027<br>(0.035) | 0.054<br>(0.049) | 0.104<br>(0.13) |
|  |  | N1 | -0.094<br>(0.065) | -0.051<br>(0.066) | -0.099<br>(0.071) | -0.108<br>(0.089) | -0.081<br>(0.045) | -0.036<br>(0.059) | -0.082<br>(0.035) | -0.131<br>(0.112) |
|  |  | P2 | 0.008<br>(0.039) | 0.062<br>(0.083) | 0.018<br>(0.043) | 0.026<br>(0.034) | 0.018<br>(0.038) | 0.037<br>(0.073) | 0.014<br>(0.035) | 0.054<br>(0.115) |
|  |  | N2 | -0.014<br>(0.023) | -0.027<br>(0.03) | -0.013<br>(0.027) | -0.011<br>(0.021) | -0.014<br>(0.041) | -0.024<br>(0.054) | -0.038<br>(0.034) | -0.074<br>(0.105) |
|  |  | P3 | 0.025<br>(0.019) | 0.015<br>(0.015) | 0.023<br>(0.019) | 0.019<br>(0.026) | 0.024<br>(0.023) | 0.043<br>(0.061) | 0.047<br>(0.037) | 0.093<br>(0.13) |
|  | Latency in<br>[s] | P1 | 0.026<br>(0.006) | 0.031<br>(0.009) | 0.029<br>(0.008) | 0.031<br>(0.007) | 0.026<br>(0.009) | 0.043<br>(0.015) | 0.038<br>(0.014) | 0.039<br>(0.01) |
|  |  | N1 | 0.048<br>(0.013) | 0.045<br>(0.013) | 0.049<br>(0.013) | 0.053<br>(0.009) | 0.049<br>(0.014) | 0.071<br>(0.025) | 0.066<br>(0.028) | 0.067<br>(0.026) |
|  |  | P2 | 0.07<br>(0.018) | 0.062<br>(0.019) | 0.077<br>(0.022) | 0.079<br>(0.013) | 0.076<br>(0.021) | 0.102<br>(0.025) | 0.092<br>(0.031) | 0.088<br>(0.028) |
|  |  | N2 | 0.088<br>(0.016) | 0.085<br>(0.033) | 0.095<br>(0.021) | 0.107<br>(0.019) | 0.097<br>(0.021) | 0.119<br>(0.024) | 0.11<br>(0.033) | 0.102<br>(0.029) |
|  |  | P3 | 0.126<br>(0.022) | 0.116<br>(0.037) | 0.13<br>(0.029) | 0.132<br>(0.019) | 0.13<br>(0.022) | 0.168<br>(0.027) | 0.17<br>(0.03) | 0.142<br>(0.039) |
| Visual cortex | Amplitude in [mV] | P1 | 0.012<br>(0.018) | 0.002<br>(0.008) | 0.024<br>(0.027) | 0.035<br>(0.028) | 0.009<br>(0.014) | 0.023<br>(0.042) | 0.046<br>(0.045) | 0.057<br>(0.054) |
|  |  | N1 | -0.049<br>(0.027) | -0.027<br>(0.019) | -0.066<br>(0.04) | -0.05<br>(0.031) | -0.039<br>(0.023) | 0.002<br>(0.027) | -0.033<br>(0.037) | -0.044<br>(0.033) |
|  |  | P2 | 0.036<br>(0.043) | 0.072<br>(0.066) | 0.069<br>(0.044) | 0.105<br>(0.059) | 0.043<br>(0.036) | 0.058<br>(0.037) | 0.08<br>(0.04) | 0.107<br>(0.053) |
|  |  | N2 | 0.006<br>(0.033) | -0.029<br>(0.035) | 0.018<br>(0.051) | 0.044<br>(0.051) | 0.019<br>(0.034) | 0.025<br>(0.03) | 0.025<br>(0.041) | 0.059<br>(0.039) |
|  |  | P3 | 0.064<br>(0.035) | 0.029<br>(0.027) | 0.043<br>(0.045) | 0.044<br>(0.042) | 0.074<br>(0.037) | 0.063<br>(0.036) | 0.072<br>(0.031) | 0.087<br>(0.033) |
|  |  | N3 | -0.06<br>(0.036) | -0.016<br>(0.012) | -0.06<br>(0.036) | -0.093<br>(0.05) | -0.076<br>(0.038) | -0.012<br>(0.022) | -0.045<br>(0.023) | -0.061<br>(0.021) |
|  |  | P4 | 0.026<br>(0.013) | 0.017<br>(0.01) | 0.032<br>(0.014) | 0.049<br>(0.026) | 0.029<br>(0.015) | 0<br>(0.023) | 0.028<br>(0.026) | 0.035<br>(0.017) |
|  | Latency in [s] | P1 | 0.024<br>(0.006) | 0.027<br>(0.007) | 0.025<br>(0.005) | 0.028<br>(0.005) | 0.023<br>(0.005) | 0.036<br>(0.011) | 0.04<br>(0.009) | 0.037<br>(0.008) |
|  |  | N1 | 0.037<br>(0.009) | 0.038<br>(0.008) | 0.037<br>(0.005) | 0.038<br>(0.003) | 0.04<br>(0.007) | 0.049<br>(0.014) | 0.051<br>(0.013) | 0.048<br>(0.008) |
|  |  | P2 | 0.056<br>(0.018) | 0.053<br>(0.013) | 0.055<br>(0.011) | 0.062<br>(0.01) | 0.062<br>(0.015) | 0.079<br>(0.018) | 0.079<br>(0.028) | 0.073<br>(0.019) |
|  |  | N2 | 0.071<br>(0.022) | 0.075<br>(0.03) | 0.074<br>(0.017) | 0.083<br>(0.013) | 0.073<br>(0.016) | 0.097<br>(0.021) | 0.102<br>(0.033) | 0.093<br>(0.024) |
|  |  | P3 | 0.102<br>(0.021) | 0.097<br>(0.039) | 0.097<br>(0.019) | 0.098<br>(0.014) | 0.099<br>(0.015) | 0.129<br>(0.023) | 0.134<br>(0.036) | 0.114<br>(0.026) |
|  |  | N3 | 0.165<br>(0.014) | 0.148<br>(0.033) | 0.16<br>(0.017) | 0.161<br>(0.017) | 0.166<br>(0.011) | 0.228<br>(0.041) | 0.222<br>(0.046) | 0.203<br>(0.044) |
|  |  | P4 | 0.225<br>(0.019) | 0.214<br>(0.037) | 0.239<br>(0.024) | 0.238<br>(0.025) | 0.228<br>(0.012) | 0.263<br>(0.045) | 0.303<br>(0.048) | 0.298<br>(0.052) |

**Figure S2.**
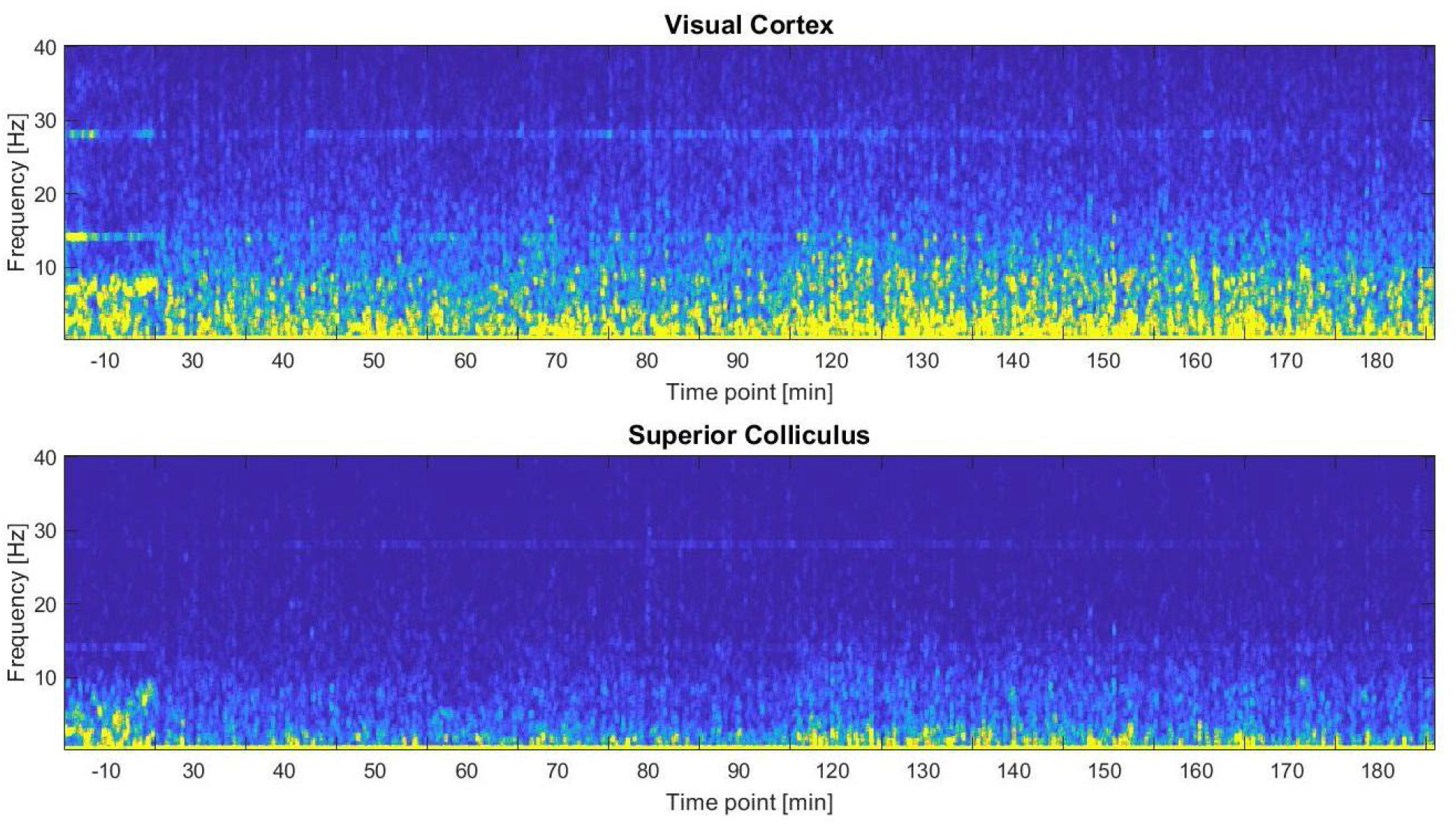
Hypnorm/midazolam anaesthesia in one animal. The spectrograms are the average of bilateral electrodes in one animal and there are hints of an SSVEP response at 14 Hz and 28 Hz. The anaesthesia was supplemented at 60 min and at 150 min.

## References

Bailey, C.J., Sanganahalli, B.G., Herman, P., Blumenfeld, H., Gjedde, A., Hyder, F., 2013. Analysis of time and space invariance of BOLD responses in the rat visual system. Cereb. Cortex 23, 210–222. https://doi.org/10.1093/cercor/bhs008

Barger, Z., Frye, C.G., Liu, D., Dan, Y., Bouchard, K.E., 2019. Robust, automated sleep scoring by a compact neural network with distributional shift correction. PLoS One 14, 1–18. https://doi.org/10.1371/journal.pone.0224642

Benjamini, Y., Yekutieli, D., 2001. The Control of the False Discovery Rate in Multiple Testing under Depency. Ann. Stat. 29, 1165–1188. https://doi.org/10.1214/aos/1013699998

Benveniste, H., Lee, H., Ding, F., Sun, Q., Al-bizri, E., Makaryus, R., Probst, S., Nedergaard, M., Stein, E.A., Lu, H., 2017. Anesthesia with Dexmedetomidine and Low-dose Isoflurane Increases Solute Transport. Anesthesiology 127, 976–988.

Desai, M., Kahn, I., Knoblich, U., Bernstein, J., Atallah, H., Yang, A., Kopell, N., Buckner, R.L., Graybiel, A.M., Moore, C.I., Boyden, E.S., 2011. Mapping brain networks in awake mice using combined optical neural control and fMRI. J. Neurophysiol. 105, 1393–1405. https://doi.org/10.1152/jn.00828.2010

Engell, A.D., Huettel, S., McCarthy, G., 2012. The fMRI BOLD signal tracks electrophysiological spectral perturbations, not event-related potentials. Neuroimage 59, 2600–2606. https://doi.org/10.1016/j.neuroimage.2011.08.079

Gao, Y.R., Ma, Y., Zhang, Q., Winder, A.T., Liang, Z., Antinori, L., Drew, P.J., Zhang, N., 2017. Time to wake up: Studying neurovascular coupling and brain-wide circuit function in the un-anesthetized animal. Neuroimage 153, 382–398. https://doi.org/10.1016/j.neuroimage.2016.11.069

Ho, J., Tumkaya, T., Aryal, S., Choi, H., Claridge-Chang, A., 2019. Moving beyond P values: data analysis with estimation graphics. Nat. Methods 16, 565–566. https://doi.org/10.1038/s41592-019-0470-3

Huneau, C., Benali, H., Chabriat, H., 2015. Investigating human neurovascular coupling using functional neuroimaging: A critical review of dynamic models. Front. Neurosci. 9, 1–12. https://doi.org/10.3389/fnins.2015.00467

Lau, C., Zhang, J.W., Xing, K.K., Zhou, I.Y., Cheung, M.M., Chan, K.C., Wu, E.X., 2011a. BOLD responses in the superior colliculus and lateral geniculate nucleus of the rat viewing an apparent motion stimulus. Neuroimage 58, 878–884. https://doi.org/10.1016/j.neuroimage.2011.06.055

Lau, C., Zhou, I.Y., Cheung, M.M., Chan, K.C., Wu, E.X., 2011b. BOLD temporal dynamics of rat superior colliculus and lateral geniculate nucleus following short duration visual stimulation. PLoS One 6. https://doi.org/10.1371/journal.pone.0018914

Lee, J.G., Hudetz, A.G., Smith, J.J., Hillard, C.J., Bosnjak, Z.J., Kampine, J.P., 1994. The Effects of Halothane and Isoflurane on Cerebrocortical Microcirculation and Autoregulation as Assessed by Laser-Doppler Flowmetry. Anesth. Analg. 79, 58–65.

Magnuson, M.E., Thompson, G.J., Pan, W., Keilholz, S.D., 2014. Time-dependent effects of isoflurane and dexmedetomidine on functional connectivity, spectral characteristics, and spatial distribution of spontaneous BOLD fluctuations. NMR Biomed. 27, 291–303. https://doi.org/10.1002/nbm.3062.Time-dependent

Niranjan, A., Christie, I.N., Solomon, S.G., Wells, J.A., Lythgoe, M.F., 2016. fMRI mapping of the visual system in the mouse brain with interleaved snapshot GE-EPI. Neuroimage 139, 337–345. https://doi.org/10.1016/j.neuroimage.2016.06.015

Østergaard, F.G., Skoven, C.S., Wade, A.R., Siebner, H.R., Laursen, B., Christensen, K.V., Dyrby, T.B., 2021. No Detectable Effect on Visual Responses Using Functional MRI in a Rodent Model of a - Synuclein Expression. eNeuro 8, 1–9. https://doi.org/https://doi.org/10.1523/ENEURO.0516-20.2021

Østergaard, F.G., Wade, A.R., Siebner, H.R., Christensen, K.V., Laursen, B., 2020. Progressive Effects of Sildenafil on Visual Processing in Rats. Neuroscience 441, 131–141. https://doi.org/10.1016/j.neuroscience.2020.06.033

Pan, W.J., Thompson, G.J., Magnuson, M.E., Jaeger, D., Keilholz, S., 2013. Infraslow LFP correlates to resting-state fMRI BOLD signals. Neuroimage 74, 288–297. https://doi.org/10.1016/j.neuroimage.2013.02.035

Pawela, C.P., Biswal, B.B., Hudetz, A.G., Schulte, M.L., Li, R., Jones, S.R., Cho, Y.R., Matloub, H.S., Hyde, J.S., 2009. A protocol for use of medetomidine anesthesia in rats for extended studies using task-induced BOLD contrast and resting-state functional connectivity. Neuroimage 46, 1137–1147. https://doi.org/10.1016/j.neuroimage.2009.03.004

Pawela, C.P., Hudetz, A.G., Ward, B.D., Schulte, M.L., Li, R., Kao, D.S., Mauck, M.C., Cho, Y.R., Neitz, J., Hyde, J.S., 2008. Modeling of region-specific fMRI BOLD neurovascular response functions in rat brain reveals residual differences that correlate with the differences in regional evoked potentials. Neuroimage 41, 525–534. https://doi.org/10.1016/j.neuroimage.2008.02.022.Modeling

Paxinos, G., Watson, C., 1998. The Rat Brain: in stereotaxic Coordinates, 4th ed. Academic Press, An imprint of Elsevier, San Diego, London.

Pradier, B., Wachsmuth, L., Nagelmann, N., Segelcke, D., Kreitz, S., Hess, A., Pogatzki-Zahn, E.M., Faber, C., 2021. Combined resting state-fMRI and calcium recordings show stable brain states for task-induced fMRI in mice under combined ISO/MED anesthesia. Neuroimage 245, 118626. https://doi.org/10.1016/j.neuroimage.2021.118626

Van Camp, N., Verhoye, M., De Zeeuw, C.I., Van der Linden, A., 2006. Light Stimulus Frequency Dependence of Activity in the Rat Visual System as Studied With High-Resolution BOLD fMRI. J. Neurophysiol. 95, 3164–3170. https://doi.org/10.1152/jn.00400.2005

Weerink, M.A.S., Struys, M.M.R.F., Hannivoort, L.N., Barends, C.R.M., Absalom, A.R., Colin, P., 2017. Clinical Pharmacokinetics and Pharmacodynamics of Dexmedetomidine. Clin. Pharmacokinet. 56, 893–913. https://doi.org/10.1007/s40262-017-0507-7

